# Benefits of commitment in hierarchical inference

**DOI:** 10.1101/658914

**Authors:** Cheng Qiu, Long Luu, Alan A. Stocker

## Abstract

Humans have the tendency to commit to a single interpretation of what has caused some observed evidence rather than considering all possible alternatives. This tendency can explain various forms of confirmation and reference biases. However, committing to a single high-level interpretation seems short-sighted and irrational, and thus it is unclear why humans seem motivated to pursue such strategy.

In a first step toward answering this question, we systematically quantified how this strategy affects estimation accuracy at the feature level in the context of two universal hierarchical inference tasks, categorical perception and causal cue combination. Using model simulations, we demonstrate that although estimation is generally impaired when conditioned on only a single high-level interpretation, the impairment is not uniform across the entire feature range. On the contrary, compared to a full inference strategy that considers all high-level interpretations, accuracy is actually better for feature values for which the probability of an incorrect categorical/structural commitment is relatively low. That is to say, if an observer is reasonably certain about the high-level interpretation of the feature, it is advantageous to condition subsequent feature inference only on that particular interpretation. We also show that this benefit of commitment is substantially amplified if late noise corrupts information processing (*e.g.*, during retention in working memory). Our results suggest that top-down inference strategies that solely rely on the most likely high-level interpretation can be favorable and at least locally outperform a full inference strategy.

## Introduction

Cognitive tasks typically require some form of statistical inference where the brain has to infer an unknown quantity based on given yet uncertain evidence and a learned statistical (generative) model of the task (Helmholtz, 1867; Jaynes, 2003; Lee and Mumford, 2003). Previous work has shown that this notion in combination with the formalism of Bayesian statistics provides an accurate description of human behavior in a broad range of tasks associated with perception (Knill and Richards, 1996), cognitive reasoning (Griffiths et al., 2010), economic decision-making (Summerfield and Tsetsos, 2012) and motor control (Wolpert, 2007). Furthermore, mental disorders such as autism and schizophrenia have been directly linked to specific computational deficiencies of this inference process (Lieder et al., 2019; Jardri and Deneve, 2013). Except for simple estimation and decision-making tasks (Ernst and Banks, 2002; Körding and Wolpert, 2004; Stocker and Simoncelli, 2006), the generative models of these tasks are hierarchical. Object recognition is a simple example of a hierarchical inference task where, at the top of the hierarchy, object categories are defined as specific distributions over some lower-level feature representation potentially across multiple levels of feature integration. Noisy observations at the lowest feature level then allow to infer the corresponding object category by inverting the hierarchical generative model (Kersten et al., 2004). Various studies have shown that human behavior in such tasks is accurately described by Bayesian statistics that fully integrate all information in the hierarchical generative model (*i.e.*, fully marginalizes) from bottom to top. These studies include models of human judgments of sameness (Van den Berg et al., 2012), of stimulus transparency (Hedges et al., 2011), or causal stimulus structure (Körding et al., 2007).

More interesting and controversial behavior has been observed in tasks that require inference at the feature rather than the top level. In these cases, the hierarchical generative model represents a hypothesis of what caused the feature, and thus ultimately the observed evidence (see Fig. 1a). Full inference dictates that in order to infer the value of the feature, an observer should consider all different possible hypotheses (*e.g.*, categorical assignments) and weigh them according to how probable they are given the observed evidence (Fig. 1b). This strategy is known as optimal model evaluation (Draper, 1995), or Bayesian model averaging (Hoeting et al., 1999), and has been previously suggested to account for human behavior in various perceptual and cognitive reasoning tasks (Duffy et al., 2006; Körding et al., 2007; Griffiths et al., 2010; Knill, 2003, 2007).

**Figure 1:**
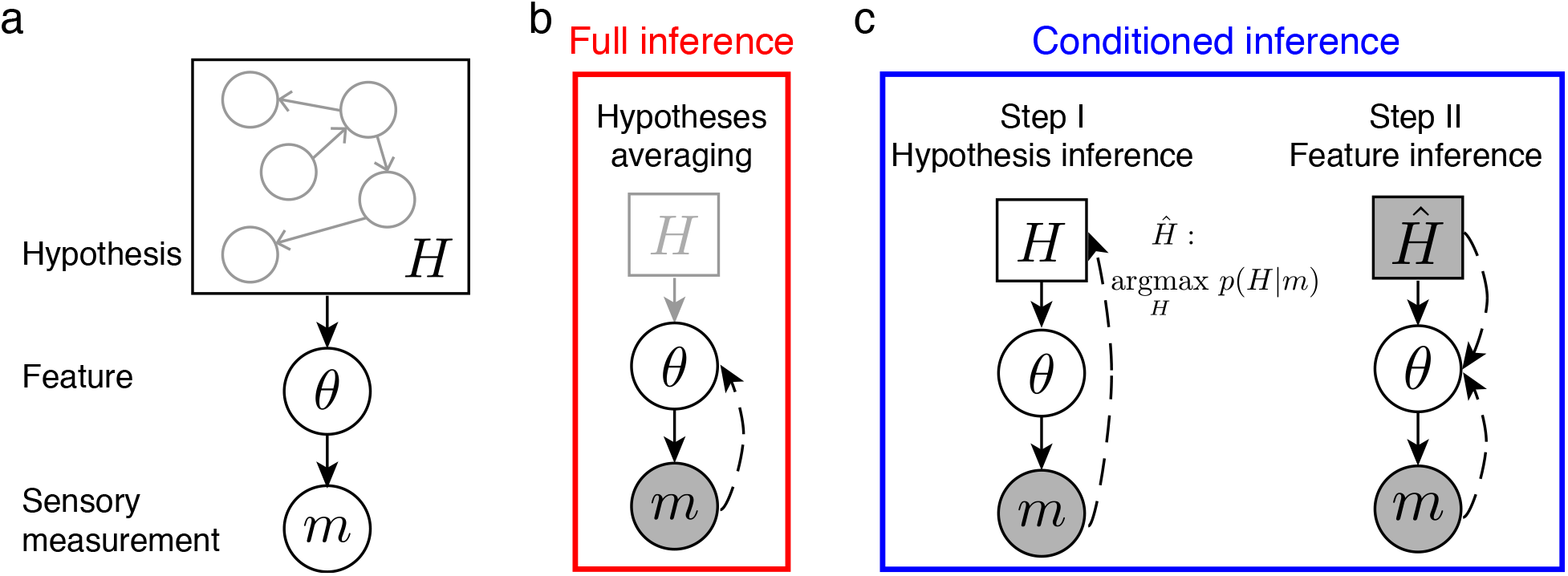
Feature inference in hierarchical generative models. (a) Graphical model that represents the generic class of hierarchical models we consider. Sensory measurement *m* is assumed to be a noisy sample of the feature value *θ*, which itself is drawn from a higher-level generative hypothesis *H*. Based on an observed value of *m* we consider two inference strategies: (b) Full inference. This strategy requires the observer to marginalize over all possible high-level hypotheses when inferring *θ*. (c) Conditioned inference. This strategy consists of two steps. First, the observer infers and commits to the most likely hypothesis *Ĥ* based on the sensory measurement *m*. Subsequently, the observer infers the feature value conditioned on the committed hypothesis *Ĥ*, rather than marginalizing over all possible hypotheses.

However, recent studies have reported conditions in which human subjects seem to abandon the averaging strategy and rather commit to a specific interpretation of what caused the evidence (model selection). For example, by making an category assignment a subject’s subsequent perceptual estimate of a low-level stimulus feature (*e.g.*, motion direction (Jazayeri and Movshon, 2007; Zamboni et al., 2016) or visual orientation (Luu and Stocker, 2018; Fritsche and de Lange, 2019)) is biased towards the assigned category on a per trial basis. These biases can be thought of as a form of consistency (Brehm, 1956) or confirmation bias (Nickerson, 1998) where the perceptual estimate aligns with and confirms the chosen category (Bronfman et al., 2015; Talluri et al., 2018). Importantly, these biases are not dependent on subjects making an explicit, overt choice of a high-level interpretation; similar biases were observed in tasks that did not require an explicit categorical choice (Wu et al., 2009; Zamboni et al., 2016; Ding et al., 2017), indicating that committing to a high-level interpretation could be an inherent inference strategy. As proposed (Stocker and Simoncelli, 2007), and refined and validated more recently (Luu and Stocker, 2018), the behavioral biases in above examples are remarkably well described by a *conditioned observer model*. The model assumes that feature inference is a sequential process: first, an observer selects the most probable hypothesis given the evidence, and then infers the feature value conditioned on the chosen hypothesis (Fig. 1c). In contrast to related models that include meta-cognitive processes with access to additional information (Fleming and Daw, 2017), the conditioned observer model assumes that the observer uses less of the available information as it discards posterior probability information by committing to only one hypothesis. Thus, the expectation is that the conditioned observer performs worse at the feature level (*i.e.*, in terms of estimation accuracy) than an observer that integrates over all possible (high-level) hypotheses.

We set out to systematically assess how strongly performance is hampered with a conditioned inference strategy. We quantitatively explored its performance compared to the full inference strategy for two simple but universal inference examples - categorical perception and causal cue combination. Using simulations over a range of generative model parameters we characterized estimation accuracy for both strategies. We show that there are conditions where the conditioned observer outperforms the full inference observer. These benefits further increase if late noise is affecting the inference process. Our results provide a useful quantitative analysis to the discussion of different inference strategies and why humans may or may not apply one and not the other.

### Inference in hierarchical generative models

We focus on the task of inferring the value of a feature *θ* based on uncertain sensory evidence *m* (measurement). We assume that *θ* is embedded in a probabilistic hierarchical generative model that can be expressed with a directed graph as shown in Fig. 1a. The hierarchical component of the model “above *θ*” represents a high-level generative hypothesis *H* about the potential values of *θ*. A simple example for such a hypothesis is the association of *θ* with a particular category. More elaborate high-level hypotheses may include different structural assumptions (*i.e.*, different graphs) that capture different contextual or causal dependencies (Körding et al., 2007; Kemp and Tenenbaum, 2008; Battaglia et al., 2013). However, our analysis is agnostic to the specific form of *H* as it only assumes that the feature *θ* is at the “bottom” of the hierarchy, and that sensory evidence *m* only directly depends on *θ* and not the rest of the hierarchy.

We consider two different inference strategies. The first strategy is to make use of the full posterior probability over hypotheses when inferring the feature (here referred to as “full inference”), that is, to marginalize over all possible high-level hypotheses (Fig. 1b). Thus, we can express the posterior over *θ* as a weighted sum

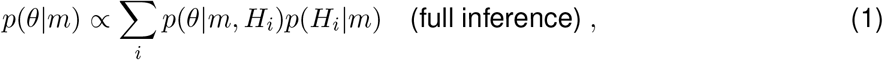

with weights given by the posterior probability of each hypothesis given the evidence, *p*(*H*_*i*_|*m*).

The second, alternative strategy represents a sequential inference process (Stocker and Simoncelli, 2007; Luu and Stocker, 2018): First, a hypothesis *Ĥ* is selected according to the posterior probability *p*(*H|m*) (here, the hypothesis with maximal posterior probability), and then the posterior over *θ* is computed conditioned on the observed evidence *m and* the chosen hypothesis *Ĥ* (Fig. 1c). The chosen hypothesis imposes a conditioned prior *p*(*θ*|*Ĥ*) that, unlike in the full inference strategy, depends on the sensory evidence and thus is potentially different for each *m* (*i.e.*, on each trial). Accordingly, the posterior probability for this second strategy can be written as

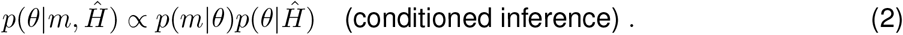

In the following we compare the performance of these two strategies in estimating *θ*. With posterior probabilities Eqs. (1) and (2) and assuming a quadratic loss function (*L*_2_-norm), we derive optimal estimators for the feature value 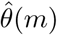 under each inference strategy, and compare their relative expected estimation error for the chosen loss function (see Appendix for results with other loss functions). We apply this error analysis to two well-known examples of hierarchical inference, categorical perception and causal cue combination systematically investigate the relative performance of the two strategies for different levels of sensory uncertainty and additional (late) processing noise.

### Example 1: Categorical perception

With *categorical perception* we refer to the task of estimating the value of a low-level (scalar) feature that is associated with multiple high-level categories^1^. A broad range of well-studied perceptual tasks and experiments represent forms of categorical perception. Categorical perception assumes the simplest hierarchical generative model where the hypothesis *H* is represented by a single node *C* reflecting the categorical assignment of feature *θ* (Fig. 2a). The generative process first involves the selection of a category *C* based on a categorical prior *p*(*C*); for simplicity, we consider here only two possible categories *C* ∈ {*C*_1_, *C*_2_}. A feature value *θ* is then sampled from the categorical feature prior *p*(*θ|C*). Finally, sensory evidence *m* is sampled from the conditional probability *p*(*m|θ*). Also, we explicitly allow the possibility that late noise may deteriorate sensory evidence *m* due to *e.g.* retention in working memory. We refer to the deteriorated sensory evidence as *m^∗^* distributed according to *p*(*m*^∗^|*m*). The specific description of the priors and conditional probabilities is provided in Fig. 2 and its caption.

**Figure 2:**
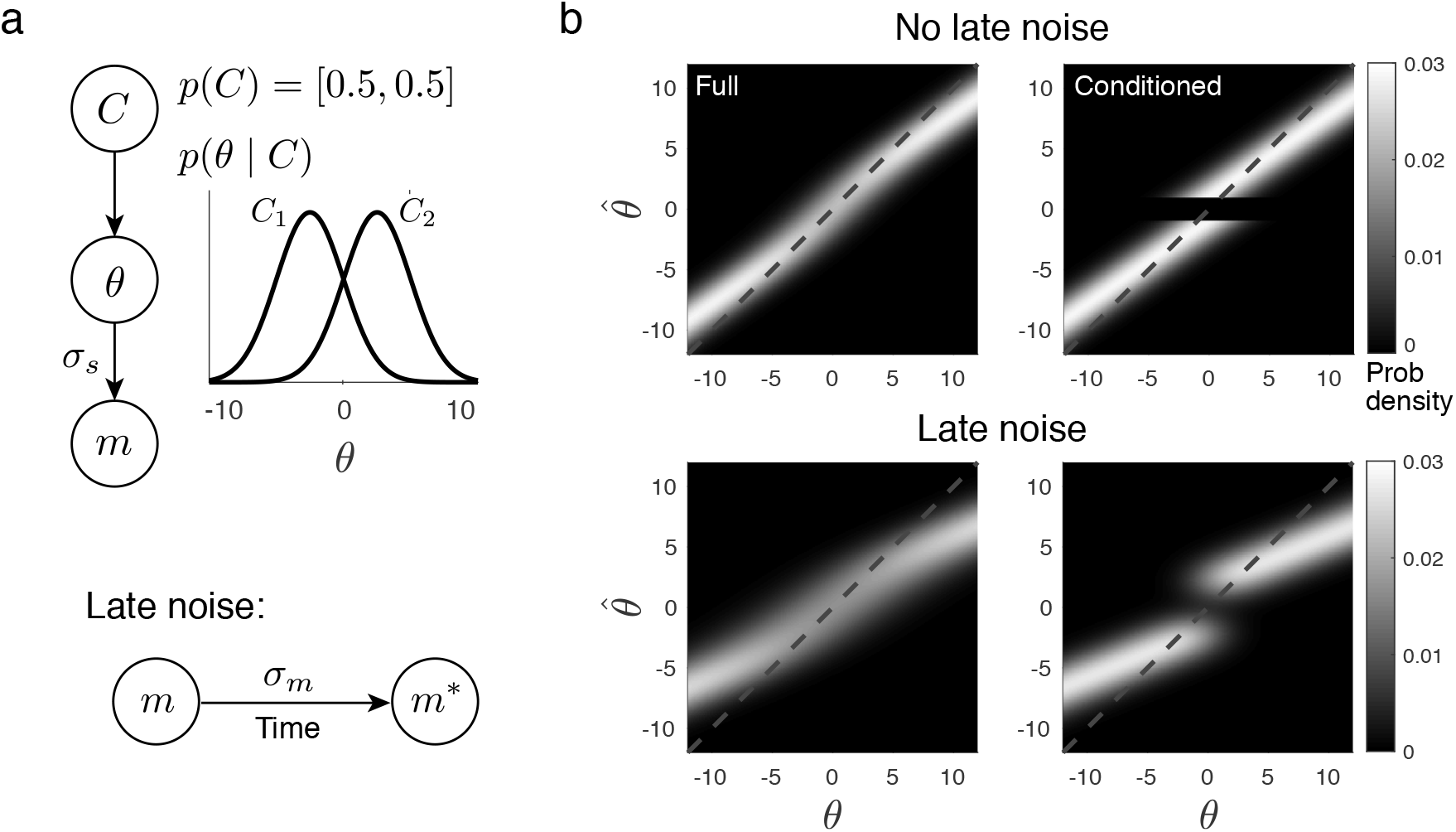
Hierarchical generative model of categorical perception, and the predicted estimate distributions for both inference strategies. (a) Generative model. For reasons of simplicity we make the following assumptions: Feature value *θ* belongs to one of two categories *C* ∈ {*C*_1_, *C*_2_} with balanced prior probabilities *p*(*C*_1,2_) = 0.5; Categorical feature priors *p*(*θ*|*C*_*i*_) are assumed to be Gaussians with identical standard deviations but different means, 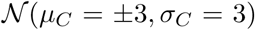; Sensory evidence *m* is a noisy sample of *θ* according to additive Gaussian noise with standard deviation *σ*_*s*_. Late additive Gaussian noise (*σ*_*m*_) may further deteriorate the sensory measurement *m* leading to *m*^∗^. (b) Estimate distributions according to either full (left) or conditioned inference (right). In each panel, vertical cross-sections represent the estimate distribution 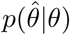. Distributions show a characteristic bimodal pattern across the category boundary for the conditioned inference strategy. Top panels show distributions without late noise (*σ*_*s*_ = 2, *σ*_*m*_ = 0); bottom panels show distributions with late noise (*σ*_*s*_ = 2, *σ*_*m*_ = 3). Units of the feature are arbitrary throughout the paper as our analysis is not limited to a specific feature.

#### Estimate distributions

Given the generative model, we can now express an optimal estimate 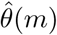 of the feature value for both inference strategies. The *full inference* strategy (Fig. 1b) marginalizes over all possible categories, resulting in the posterior distribution

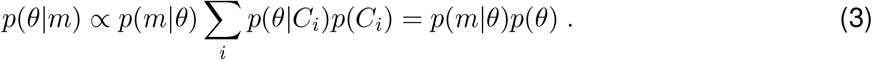

Minimizing mean squared-error (*i.e.*, minimizing *L*_2_ loss) we find the optimal estimator according to the full inference strategy as

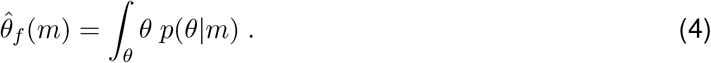

In contrast, the *conditioned inference* strategy consists of two inference steps (Fig. 1c). First, the most probable category is chosen based on the sensory evidence *m* according to

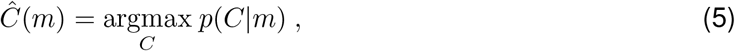

where the posterior is defined as *p*(*C|m*) ∝ *p*(*C*)*∫*_*θ*_ *p*(*m|θ*)*p*(*θ|C*). Second, the posterior over *θ* is computed conditioned on the chosen category *Ĉ*(*m*), thus

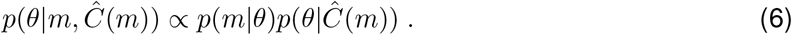

Finally, the optimal estimator is given as

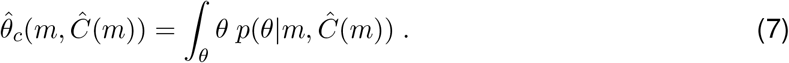

Note, that for both strategies the optimal estimator represents a monotonic mapping between the evidence and the estimate. Thus, we can obtain the estimate distributions 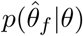 and 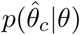 with a change of variable for the measurement distribution *p*(*m|θ*), replacing *m* with the estimate 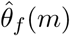 and 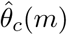 according to Eqs. (4) and (7), respectively. Figure 2b shows the resulting distributions for both strategies given the specific parameter settings of our generative model. Note that the estimate distributions fundamentally differ; the conditioned inference strategy exhibits a characteristic bimodal distribution for *θ* values close to the category boundary, which matches a range of experimental results (Jazayeri and Movshon, 2007; Zamboni et al., 2016; Luu and Stocker, 2018).

Consider the possibility that late noise may further deteriorate sensory evidence *m* (*e.g.* due to retention in working memory (Schneegans and Bays, 2018)) we update the formulations of the optimal estimators accordingly. Assuming that late noise is additive Gaussian (with *σ*_*m*_), we compute the optimal estimate and the estimate distribution for the full inference strategy as discussed above (Eqs. (3) and (4)) but replacing *p*(*m|θ*) with *p*(*m*^∗^|*θ*) = *∫*_*m*_ *p*(*m*^∗^|*m*)*p*(*m|θ*) where *m*^∗^represents the corrupted sensory evidence. For the conditioned inference strategy, we assume that late noise affects feature estimation only but not the selection of the category in step I. Thus, the computation of *Ĉ* is as above (Eq. (5)). The optimal feature estimator, however, changes to

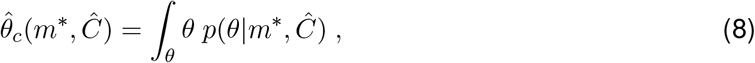

where *p*(*θ*|*m*^∗^, *Ĉ*) ∝ *p*(*m*^∗^|*θ*)*p*(*θ*|*Ĉ*). By a change of variable in *p*(*m*^∗^|*θ*, *Ĉ*) substituting *m^∗^* with the estimate 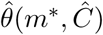, we find 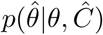. Then the estimate distributions are obtained as

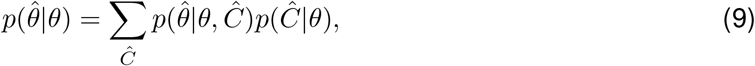

where *p*(*Ĉ*|*θ*) = ∫_*m*_*p*(*Ĉ*|*m*)*p*(*m*|*θ*). Figure 2b shows the estimate distributions given the feature value for full and conditioned inference for moderate level of late noise.

#### Relative accuracy of full vs. conditioned inference strategy

Having defined the optimal estimate distributions for both the full and the conditioned inference strategy, we next perform a systematic quantitative comparison between the estimation accuracy for various model parameters and categorical priors. We defined relative accuracy as the ratio between the expected estimation errors for both inference strategies. Because the optimal estimators were derived with regard to a quadratic loss function (*L*_2_-norm), we accordingly defined the expected estimation error as the mean squared-error (MSE). We computed relative accuracy both *locally*, *i.e.*, for both strategies we computed the MSE for each *θ* separately as 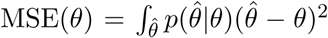 and then took the ratio, as well as *globally*, *i.e.*, we marginalized the local error over the total distribution of feature values MSE = *∫*_*θ*_ MSE(*θ*)*p*(*θ*) and then took the ratio.

We considered two general conditions of categorical feature priors. First, we assumed the priors *p*(*θ|C*_1,2_) to be overlapping Gaussian distributions (Fig. 3a). Local relative accuracy with and without late noise is shown in Fig. 3b. Both curves show a similar, characteristic shape. For *θ* values close to the category boundary, the full inference strategy provides better estimates. However, for *θ* values that are farther away than a certain distance from the category boundary, the conditioned inference strategy consistently outperforms the full inference strategy. This came as a surprise. The advantage of the conditioned inference strategy is amplified if we further assume that late noise (*σ*_*m*_) corrupts the inference process. The distance from the category boundary at which the conditioned inference strategy starts to provide superior estimates corresponds to a certain probability level of making the correct categorical assignment *Ĉ*, *i.e.*, a certain level of the psychometric function shown in Fig. 3c. It demonstrates that for conditions where the observer is reasonably certain of the correct feature category (*e.g.*, a probability correct of *p* = 0.92 if there is no late noise) the conditioned inference strategy outperforms the full inference strategy. As expected, global relative accuracy for the conditioned inference strategy is always inferior to the full inference strategy if there is no late noise (Fig. 3d). This is because the benefits of the full inference strategy for *θ* values around the category boundary make up for the deficits for *θ* values outside of that range. The situation changes, however, if substantial late noise impacts the inference process. In this case the conditioned strategy can also globally outperform the full inference strategy (Fig. 3d).

**Figure 3:**
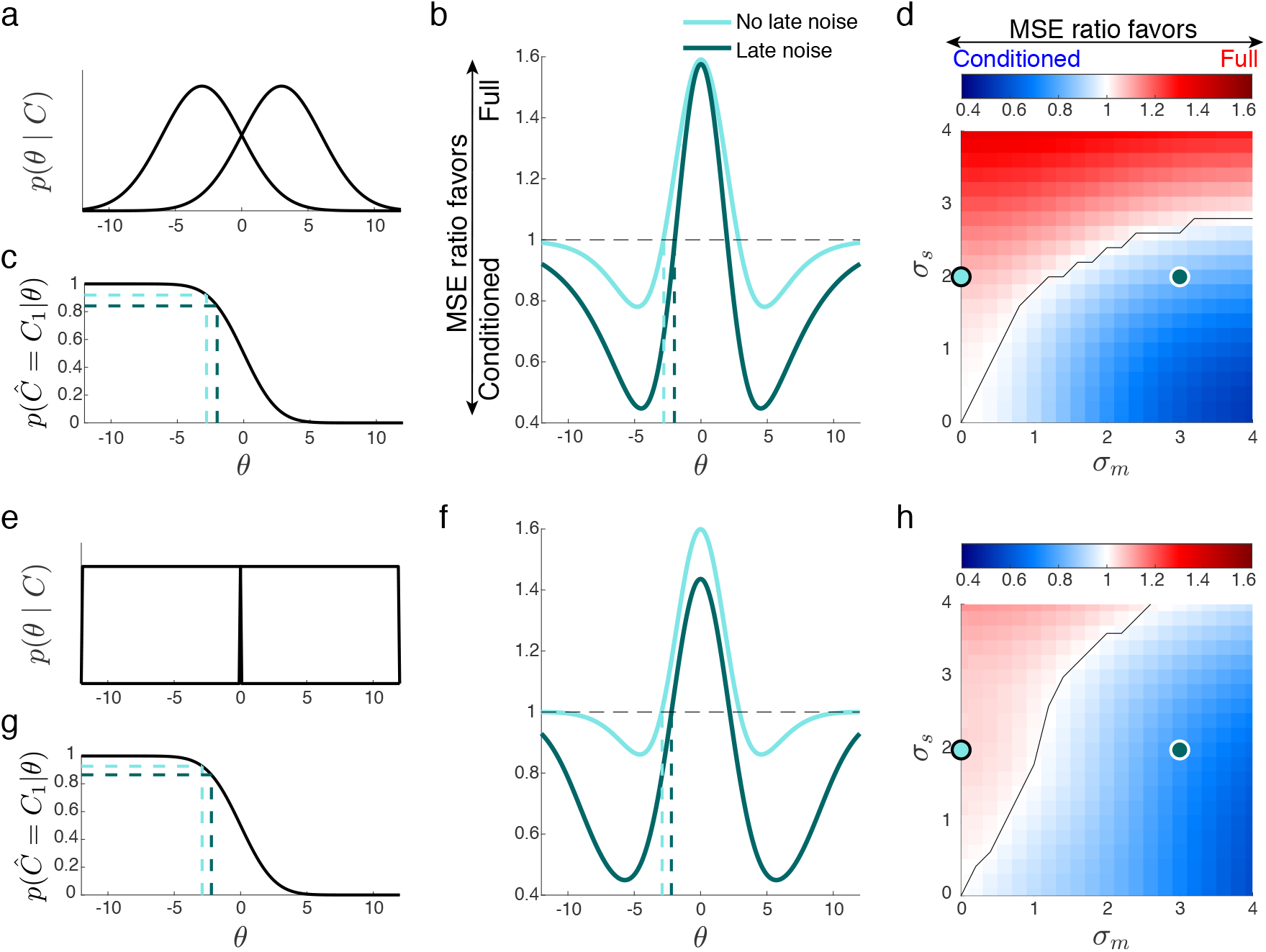
Relative estimation accuracy for conditioned and full inference in categorical perception. (a) Overlapping categorical feature priors (*µ*_*C*_ = *±*3, *σ*_*C*_ = 3). The categorical prior *p*(*C*) is symmetric. (b) Relative accuracy (MSE(*θ*) ratio) for the two strategies as a function of *θ* with sensory noise *σ*_*c*_ = 2 and late noise either absent (light color) or present (*σ*_*m*_ = 3, dark color). The distance from the category boundary at which the conditioned inference strategy starts to provide superior estimates corresponds to a probability of *p* = 0.92 for the model observer making a categorical assignment *Ĉ* (*p* = 0.85 if late noise is present; dashed lines - see also (c)). (c) Psychometric curve of the model observer indicating the probability of assigning *θ* to category *C*_1_. (d) Global MSE ratio as a function of *σ*_*s*_ and *σ*_*m*_. Blue shades represent conditions for which conditioned inference outperforms full inference, red shades indicate the opposite. The black curve labels the conditions for which both strategies lead to equal performance. The two dots correspond to the two conditions shown in (b). (e-h) Prior distributions, psychometric curve, local and global relative accuracy when the two categorical feature priors are box-shaped and non-overlapping.

We obtain similar results for non-overlapping categorical feature priors (Fig. 3e). This corresponds to many of the estimation tasks reported in previous studies (Jazayeri and Movshon, 2007; Zamboni et al., 2016; Luu and Stocker, 2018). Local relative accuracy shows very similar curves (Fig. 3f), indicating the supremacy of the conditioned inference strategy for *θ* values that correspond to similar certainty levels in the categorical assignment (Fig. 3g). Also the global accuracy comparison shows similar behavior (Fig. 3h). While we limited our accuracy analysis in the main paper for the case of minimizing squared-error (*i.e.*, the *L*_2_ norm), we explored other commonly used error metrics as well. We found that the above results of our analysis qualitatively generalize (see Appendix).

### Example 2: Causal cue combination

Our second example is often referred to as causal cue combination (Körding et al., 2007). When tasked to estimate the unknown value of a feature, human observers correctly combine different sensory cues if the cues are in sufficient agreement with the interpretation that they originate from the same feature value (Jacobs, 1999; Ernst and Banks, 2002; Alais and Burr, 2004; Fetsch et al., 2009; Butler et al., 2010). However, human observers do not integrate cues that are inconsistent with such interpretation (cue conflict) and thus signal different underlying feature values (Roach et al., 2006; Wallace et al., 2004).

This integration versus segregation distinction can be modeled within the generative hierarchical framework we discuss here (Fig. 1a). However, different to the categorical perception example, the hypotheses now represent two different causal structures (Fig. 4a): Either the cues *m*_1,2_ represent evidence of a single feature *θ* and thus should be combined (common cause hypothesis), or they represent two different features *θ*_1,2_, in which case they should be treated independently (independent causes hypothesis). The full inference strategy for estimating the feature values considers both structural hypotheses *S* ∈ {*S*_1_, *S*_2_} according to their posterior probabilities when inferring the posterior density over the features *p*(*θ*_1_*, θ*_2_*|m*_1_*, m*_2_). In contrast, the conditioned inference strategy first commits to an interpretation *Ŝ* of the most probable structural hypothesis based on the observed sensory evidence, and then computes the posterior density over the feature values conditioned only on the chosen structure, thus *p*(*θ*_1_,*θ*_2_|*m*_1_*m*_2_,*Ŝ*).

**Figure 4:**
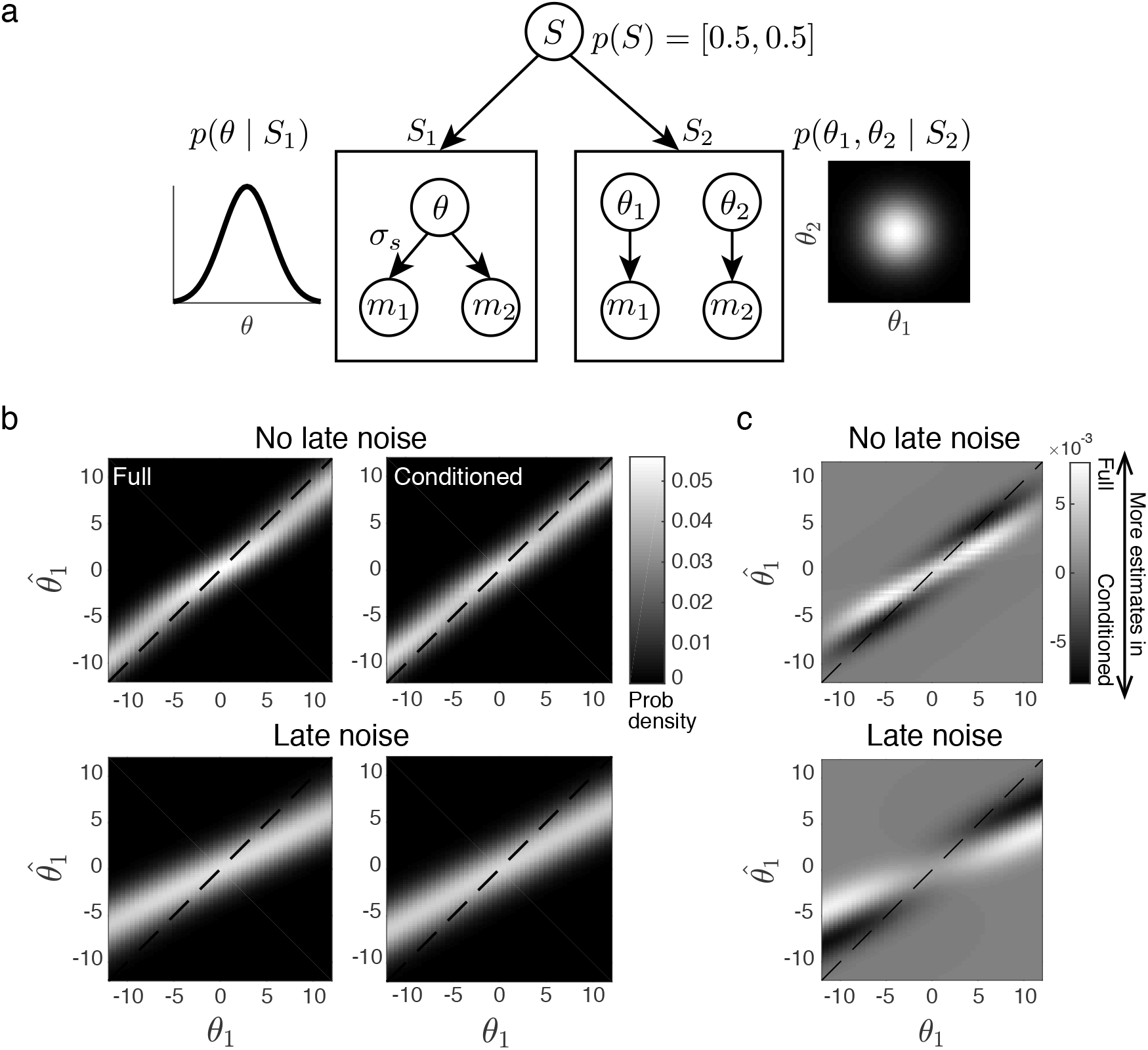
Causal cue combination. (a) Graphical model reflecting two possible generative structures (hypotheses): the observed sensory measurements *m*_1,2_ may be generated by a common source (*S*_1_) or two independent sources (*S*_2_). For reasons of simplicity we assume the structure prior *p*(*S*) to be equal, and the prior over feature values to be normally distributed: 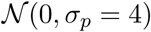 for *S*_1_, and 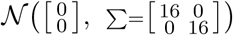 for *S*_2_. Furthermore, we also assume the observed sensory measurements to be independent samples drawn from Gaussian distributions 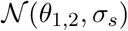. (b) Estimate distributions for the full and the conditioned inference strategy, either without (top row) or with late noise (bottom row; Gaussian 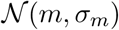. Each vertical cross-section represents the estimate density 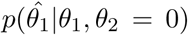 with *σ*_*s*_ = 2 and late noise *σ*_*m*_ either 0 or 3. (c) Difference of the estimate distributions in (b). For *θ*_1_ values substantially different from *θ*_2_ = 0, conditioned inference produces more veridical estimates than full inference.

#### Feature estimation

According to the hierarchical generative model shown in Fig. 4a we can define the optimal estimator under each inference strategy. For *S* = *S*_1_ (common cause), the corresponding posterior distribution at the feature level is

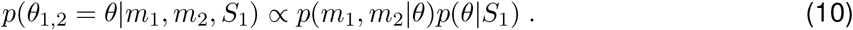

For *S* = *S*_2_ (independent causes), the posterior distribution changes to

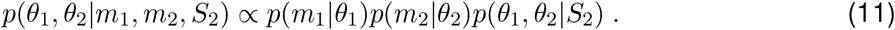

With a full inference strategy, we compute the total posterior as the average posterior under both hypotheses (Eqs. (10) and (11)) weighted by the posterior probability of each hypothesis, thus

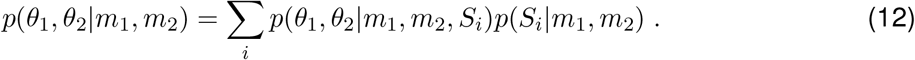

The posterior *p*(*S|m*_1_, *m*_2_) is different for the two hypothesis, namely

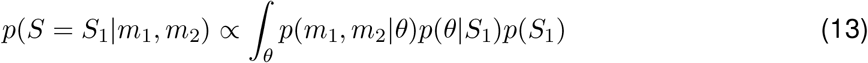

and

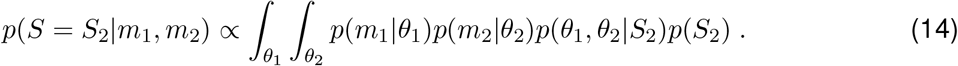

As in our first example, we define the optimal estimate of the feature vector [*θ*_1_, *θ*_2_] with regard to the *L*_2_-norm which corresponds to the expectation over the total posterior Eq. (12) (mean of posterior), thus

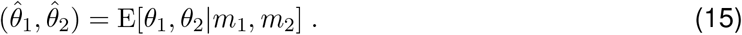

With the conditioned inference strategy we first infer the most probable causal structure,

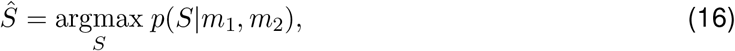

with *p*(*S|m*_1_, *m*_2_) as defined above (Eqs. (13) and (14)). The optimal estimate of the feature vector is again the expectation over the posterior distributions; however, now conditioned on *Ŝ* as well, thus

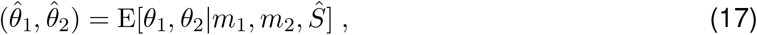

where the posterior is given as in Eqs. (10) and (11), according to *Ŝ*.

For both strategies, the estimate distributions, 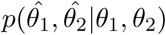, are obtained by marginalizing over the measurement distributions *p*(*m*_1_, *m*_2_|*θ*_1_, *θ*_2_) (change of variable). Figure 4b shows the estimate distributions for 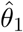 for both strategies with setting *θ*_2_ = 0, equal structural priors *p*(*S* = *S*_1,2_) = 0.5, and Gaussian distributed feature values *θ*_1,2_ and measurement distributions (see figure/caption for details). Distributions with and without late noise are shown where late noise was again assumed to be additive, independent Gaussian noise.

Unlike in the categorical perception example, the resulting estimate distributions for both inference strategies are very similar given the chosen model parameters. Plotting the difference in distribution density (Fig. 4c), however, shows that the estimates are more veridical for the full inference strategy for values of *θ*_1_ close to *θ*_2_, whereas the conditioned strategy leads to more veridical estimates for conditions where there is a substantial cue conflict, *i.e.*, *θ*_1_ ≠ *θ*_2_.

#### Comparing estimation accuracy of full vs. conditioned inference strategy

Similar to the previous example, we analyzed the relative estimation accuracy of the two inference strategies by computing both the local MSE(*θ*_1_, *θ*_2_) as well as the global MSE (integral over all *θ*_1,2_). Results are shown in Fig. 5. For feature value pairs (*θ*_1_, *θ*_2_) that support both the common and the independent source hypothesis, full inference outperforms conditioned inference (Fig. 5c). For feature values, however, that clearly favor one structural interpretation over the other, the conditioned inference strategy is beneficial. The relative accuracy comparison is similar to the categorical perception example as shown in Fig. 5d. Again, in situations where observers are reasonably certain about the high-level interpretation, conditioned inference proves to be the better strategy. This advantage is further amplified if late noise affects feature inference (Fig. 5e).

**Figure 5:**
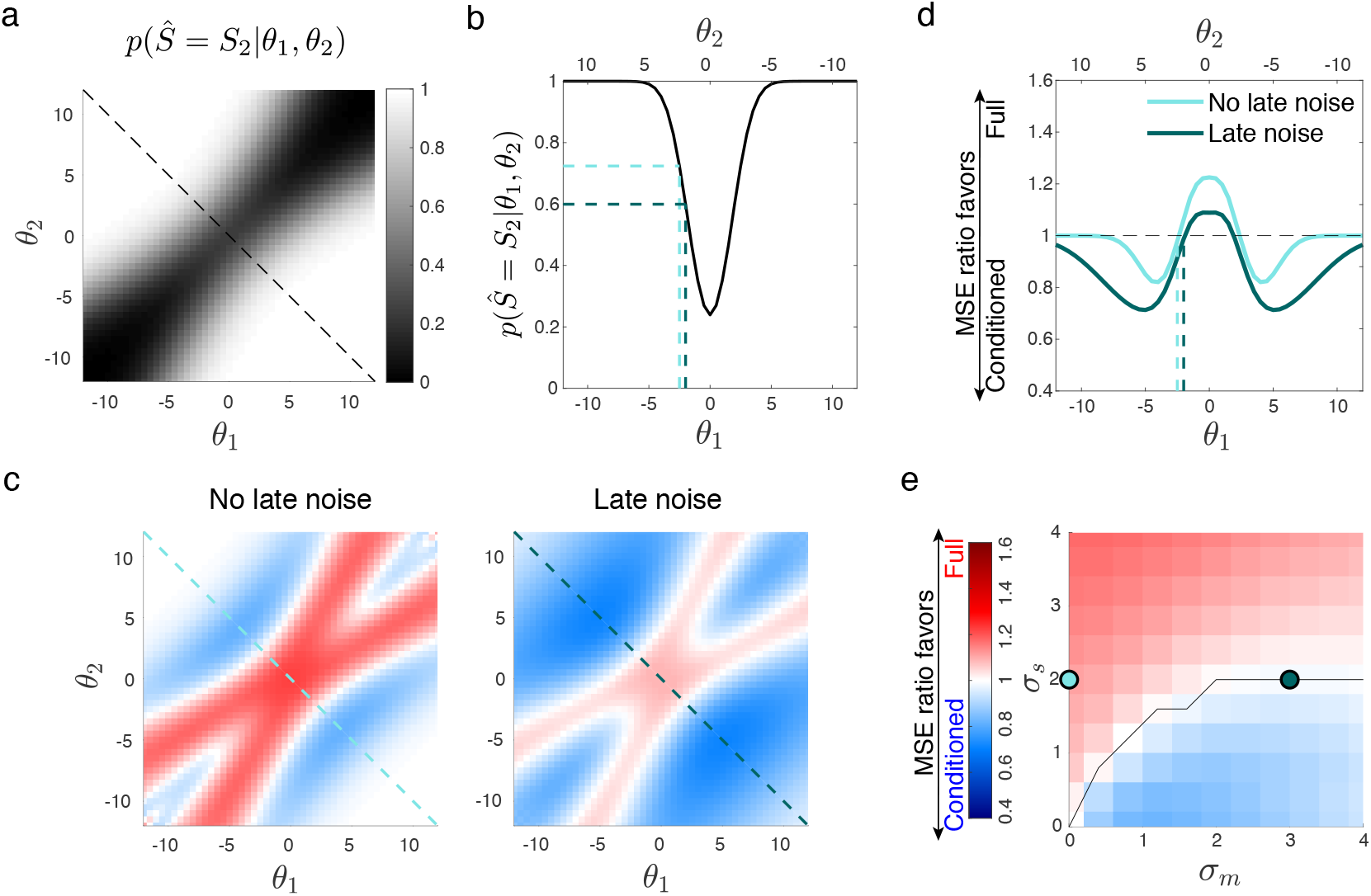
Relative performance between conditioned and full inference for causal cue combination. (a) Probability of inferring a structure with independent sources (*S*_2_) as a function of feature values. When *θ*_1_ and *θ*_2_ are far apart, they most likely represent independent sources, whereas when they are close, a common source is more plausible. (b) Cross-section through the probability density surface in (a) along the off-diagonal (dashed line). (c) Relative performance in estimating *θ*_1_ and *θ*_2_ (ratio of net MSE) for sensory noise *σ*_*s*_ = 2 and no (*σ_m_* = 0) or moderate late noise (*σ*_*m*_ = 3). Blue indicates feature values for which conditioned inference outperforms full inference. These values correspond to a regime where decisions about *Ŝ* are relatively certain as shown in (a). Red indicates feature values for which full inference is better. With late noise, the benefits of conditioned inference are amplified. (d) Local relative MSE along the off-diagonal *θ*_1_ = *−θ*_2_ (dashed lines in (c)). Conditioned inference outperforms full inference when the observer is relatively certain that *θ*_1_ and *θ*_2_ come from independent sources (dashed lines in (b)). (e) Global MSE ratio as a function of sensory (*σ*_*s*_) and late noise (*σ*_*m*_).

## Discussion

Since first proposed (Stocker and Simoncelli, 2007), data from a growing number of psychophysical studies suggest that humans perform conditioned inference in certain situations (Jazayeri and Movshon, 2007; Zamboni et al., 2016; Wu et al., 2009; Ding et al., 2017; Fritsche and de Lange, 2019; Luu and Stocker, 2018); *i.e.*, they commit to a single high-level interpretation rather than taking into account all possible high-level interpretations when performing low-level feature inference. This also matches recent results showing that post-decision confidence reports overemphasize information supporting a decision (Peters et al., 2017). In this paper, we presented an extensive quantitative analysis of how a conditioned inference strategy affects the accuracy with which an observer is able to estimate low-level features. Using model simulations, we show that considering all possible high-level interpretations does not necessarily provide better accuracy across the entire feature range. Specifically, under conditions where the observer is reasonably certain about the high-level interpretation, committing to a single interpretation is better. Although our conclusions must necessarily remain limited to the specific examples we have analyzed, categorical perception and causal cue combination, the results for these two well-known yet quite different inference problems are qualitatively very similar and robust to variations in model parameters (such as *e.g.* different prior distributions - see Appendix). This suggest that these *benefits of commitment* may generally apply to the type of hierarchical problems in perception and cognition we consider here (Fig. 1a).

We also demonstrated that a conditioned inference strategy becomes increasingly favorable when late noise is affecting the inference process, to a point where it can globally outperform a full inference strategy. This has implications for models and theories of working memory formation and recall, and in general, for inference tasks that evolve over time (Gold and Stocker, 2017). Specifically, it suggests that the commitment to a high-level interpretation can be seen as creating a robust summary representation of sensory evidence, which then, at the time of recall, is used in order to improve inference accuracy. In fact, conditioned inference can predict some of the memory biases that have been reported in the recall of color (Bae et al., 2015) or other low-level visual features (Ding et al., 2017; Luu et al., 2017). It provides a normative motivation for memory biases and potentially other similar forms of confirmation biases (Talluri et al., 2018; Lange et al., 2019).

Our results also have theoretical implications. While a full inference strategy provides globally better estimation accuracy, a conditioned inference strategy is more accurate in situations where the observer can be reasonably certain that the high-level interpretation is correct. While this certainty level does not correspond to a fixed probability value and depends on the specifics of the generative model, we found that it lies within a relative limited range for a given generative model (see dashed lines in Fig. 3). This suggests that a mixed inference strategy that uses either conditioned or full inference depending on the certainty in the high-level interpretation could globally outperform full inference under any condition. It would require a meta-cognitive mechanism that monitors and evaluates the observer’s certainty in all possible high-level hypotheses. We are not aware of any previous proposal of such a “supra-optimal” mixed inference strategy. However, it would provide a quantitative argument for why meta-cognitive processes that monitor decision confidence may be useful. Future work will show if human inference behavior shows any evidence for such a mixed strategy. Our results also provide as a useful quantitative analysis for the ongoing general debate about optimality/suboptimality in human perceptual and cognitive behavior (Rahnev and Denison, 2018; Stocker, 2019).

An obvious but willful limitation of our study is that we only analyzed accuracy at the feature level (*i.e.*, a loss function only including *θ*). However, cognition involves simultaneous and co-joint inference processes at all hierarchical levels that together form a holistic representation. Thus, it is likely that human inference strategies have evolved with regard to error metrics that are defined at the level of such holistic representations. For example, in the case of categorical perception (see Fig. 2a), the error metric would not only include errors at the feature but simultaneously also at the category level. Furthermore, one can easily make the case that such metric includes certain measures of consistency across the hierarchy; errors at the feature level would be irrelevant if at the same time the category assignment of the feature is already wrong (“why bother about error at the low level if inference is wrong at the top level?”). It is easy to show that conditioned inference globally outperforms full inference if the analysis is limited to those trials where the observer is correct at the category level: As both strategies have equal performance in getting the category right, conditioned inference is always beneficial because it uses the more specific feature prior 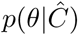 where 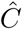 is correct. Also, we have recently shown how conditioned inference avoids inconsistencies of the representations across the hierarchy under conditions of late noise (Luu and Stocker, 2018), which may be a mechanism to avoid states of cognitive dissonance (Brehm, 1956; Festinger, 1957).

Our comparison of the inference strategies did not consider potential differences in computational and resource costs. There is growing evidence that cognitive and perceptual inference processes are optimized also with regard to such costs, leading to inference strategies that are commonly referred to as bounded rationality (Simon, 1984; Gershman et al., 2015). Intuitively, conditioned inference is a simpler and computationally less costly strategy because it performs feature inference only for one high-level hypothesis. Future work will be needed to investigate in more detail the benefits (or drawbacks) of a conditioned inference strategy with regard to hierarchical error metrics and other costs such as computational complexity.

Finally, given that many inference problems are of the general hierarchical type considered here, conditioned inference strategies may be more ubiquitous than we recognize at the moment. The problem is that conditioned inference not always leads to clearly identifiable behavioral signatures such as the bimodal estimate distributions in categorical perception (Fig. 2b). In some cases behavioral effects may be small and difficult to extract from experimental data (Fig. 4b,c). Potential candidates for conditioned inference strategies are context-dependent perceptual tasks where different contextual interpretations represent the different high-level hypotheses. Examples include tilt estimation of textured surfaces with competitive priors (Knill, 2003), orientation estimation affected by center-surround integration or segmentation (Schwartz et al., 2009; Coen-Cagli et al., 2015; Qiu et al., 2013), lightness perception affected by perceived surface curvature (Knill and Kersten, 1991), or the perceived brightness of a gray patch depending on its spatial context (Adelson, 1993). A detailed quantitative modeling approach will allow us to determine the extent to which conditioned inference is a ubiquitous and general inference mechanism of the human brain.

## Acknowledgments

We thank current and past members of the Computational Perception and Cognition laboratory for their various forms of feedback and inspirations. Some of the ideas have been described in a paper posted on BioRxiv (Luu et al., 2017). This work has been supported in part by the National Science Foundation of the United States of America (CAREER award 1350786 to AAS), and in part by the University of Pennsylvania.

## A Appendix

### A.1 Categorical perception

Our analysis in the main text was limited to comparing optimal estimators under a *L*_2_-norm loss function (MSE estimates) and symmetric top-level priors. In order to get a sense of the generality of our results, we also compared the accuracy of optimal estimators formulated under both inference strategies for a *L*_1_-norm loss function (measured with regard to absolute error, correspondingly) as well as for asymmetric category priors *p*(*C*) (for *L*_2_-norm).

#### A.1.1 Minimizing *L*_1_-norm loss

Using the same generative model and model parameters as in the main text (Fig. 2a) we formulated optimal estimators for each inference strategy as the median of the posterior distributions Eqs. (3) and (6), respectively, and performed the same accuracy comparison as in the main text with regard to absolute error. As shown in Fig. A.1, the results are qualitatively very similar to the *L*_2_-norm case (Fig. 3).

**Figure A.1:**
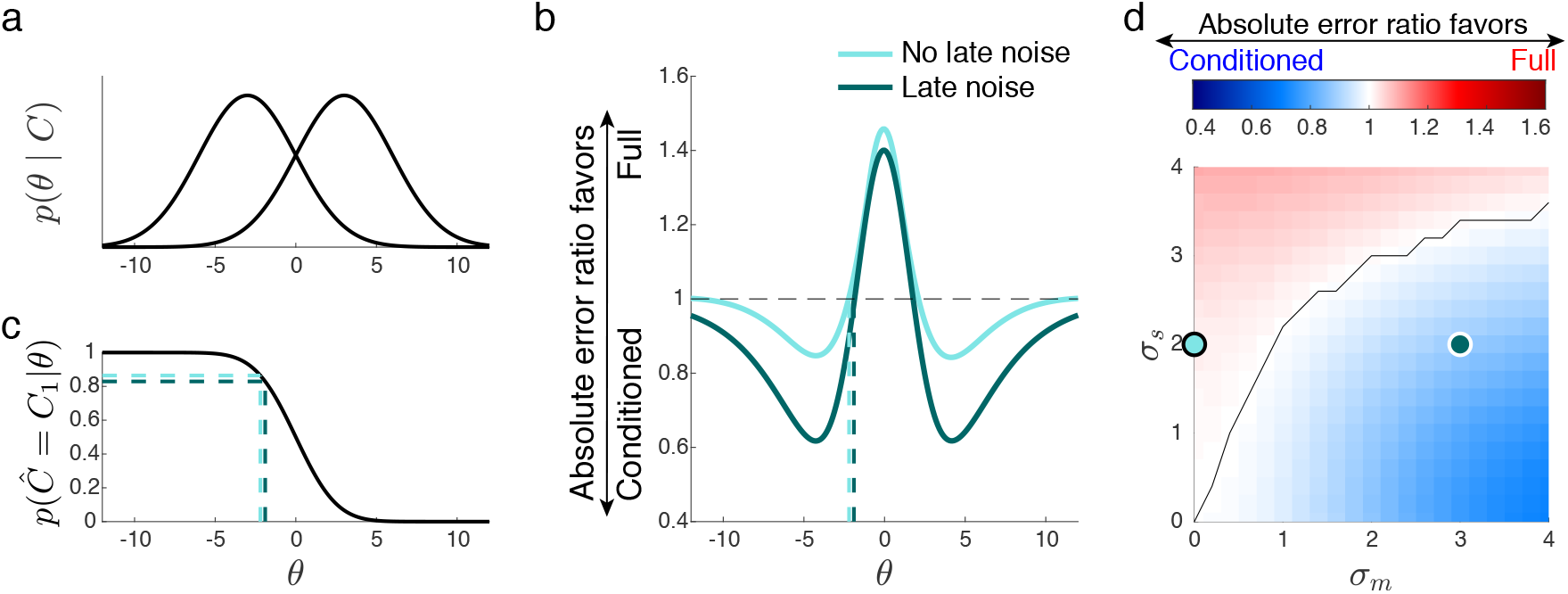
Comparison of the full and conditioned inference strategy when the observer is optimized with regard to a *L*_1_-norm loss function. Accuracy comparison is correspondingly assessed in terms of *L*_1_-norm loss (absolute error). Panels correspond to panels in Fig. 3. All other parameters were identical to the example in the main text.

#### A.1.2 Asymmetric category priors

We also explored the scenario that one category was more likely than the other, *i.e.*, the case *p*(*C*_1,2_) = [0.2, 0.8]. Figure A.2 shows the results for both overlapping and non-overlapping categories (panels are similarly organized as in Fig. 3). With unequal category priors, the psychometric curves *p*(*C|θ*) are shifted away from the categorical boundary (*e.g., p*(*C*_1_*|θ*) is shifted away from the more prevalent category *C*_2_). Compared to the equal prior case, the pattern of local relative accuracy maintains, although it is also shifted (Fig. A.2b,f). The asymmetric categorical priors exaggerate the differences between posteriors *p*(*C*_1_|*m*) and *p*(*C*_2_*|m*) making full inference act more like conditioned inference. The differences in terms of the performance between the two strategies are consequently reduced. In the extreme case where *p*(*C*_1,2_) = [0, 1] (only one category is possible), the two strategies are functionally identical.

### A.2 Causal cue combination: different feature prior

We analyzed the causal cue combination example for the case of an approximately uniform categorical feature prior (*σ_p_* = 100). The relative performance pattern is qualitatively similar compared to the example in the main text.

**Figure A.2:**
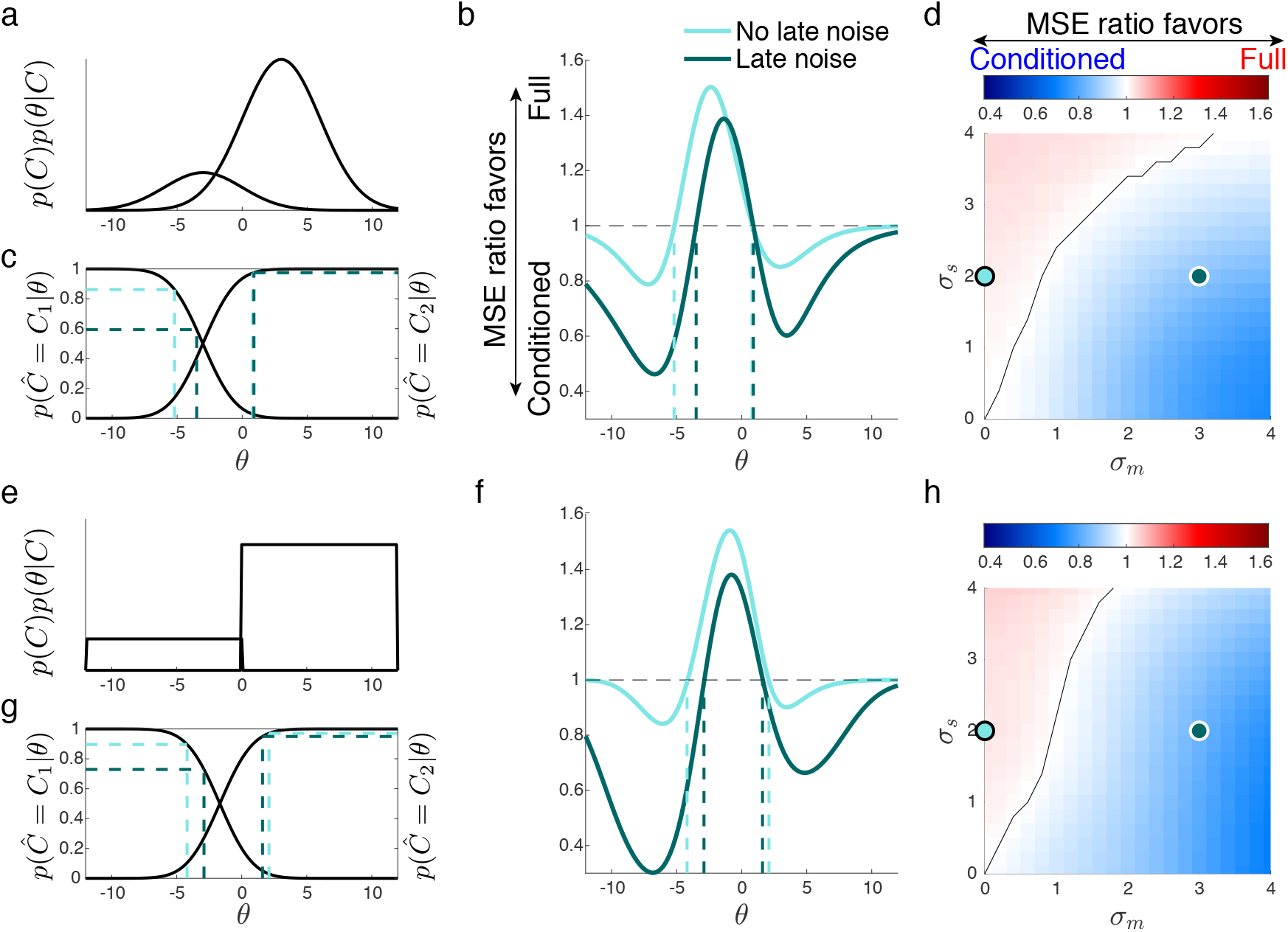
Estimation accuracy for asymmetric category priors. The panels are similarly organized and labeled as in Fig. 3. (a) Categorical feature priors multiplied with the categorical prior for illustration purposes. (b) Local performance comparison between conditioned and full inference. Larger benefits are seen when there is reasonable evidence for committing to the less prevalent category (as shown in (c)). (c) Psychometric curve indicating the probability of assigning *θ* to category *C*_1,2_. (d) Global MSE ratio as a function of sensory (*σ_s_*) and late noise (*σ*_*m*_). (e-h) Same as (a-d) for non-overlapping categorical feature priors.

**Figure A.3:**
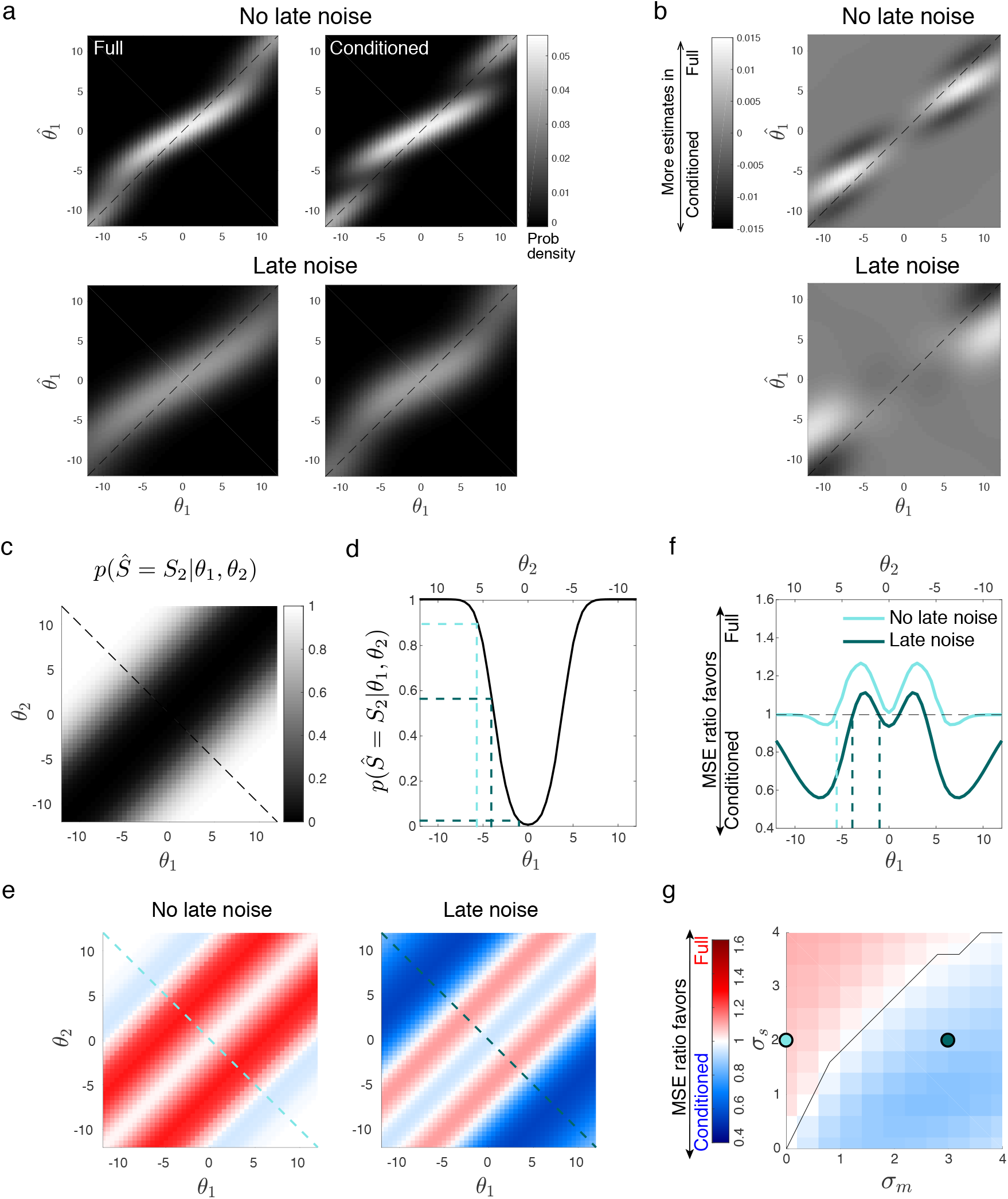
Simulations of causal cue combination with approximately uniform feature prior (*σ*_*p*_ = 100). (a,b) Estimate distributions and their differences between the two strategies. (c-g) Relative MSE accuracy. Panels correspond to panels in Fig. 5.

## Footnotes

1 This in contrast to *categorization*, which is the task of inferring the category based on evidence about the feature value.

